# Heuristics for the De Bruijn Graph Sequence Mapping Problem

**DOI:** 10.1101/2023.02.05.527069

**Authors:** Lucas B. Rocha, Said Sadique Adi, Eloi Araujo

## Abstract

In computational biology, mapping a sequence `s` onto a sequence graph `G` is a significant challenge. One possible approach to addressing this problem is to identify a walk `p` in `G` that spells a sequence which is most similar to `s`. This problem is known as the Graph Sequence Mapping Problem (`GSMP`). In this paper, we study an alternative problem formulation, namely the De Bruijn Graph Sequence Mapping Problem (`BSMP`), which can be stated as follows: given a sequence `s` and a De Bruijn graph `G`_*k*_ (where `k`≥ 2), find a walk `p` in `G`_*k*_ that spells a sequence which is most similar to `s` according to a distance metric. We present both exact algorithms and approximate distance heuristics for solving this problem, using edit distance as a criterion for measuring similarity.

## 1 Introduction

A very relevant task in computational biology is to map a sequence onto another for comparison purposes. Typically, one sequence is compared to a reference sequence, which is considered a high-quality sequence representing a set of sequences [9, 10]. On the other hand, the reference sequence is biased as it represents only a limited set of sequences and it is unable to account for all possible variations. One way to overcome this bias is to represent multiple sequences as another robust structure [6], such as the sequence graph or De Bruijn graph [7, 13, 14].

The *sequence graph* is a graph such that each node is labeled with one or more characters and the *simple sequence graph* is one where each node is labeled with exactly one character [7]. In the *De Bruijn graph* [13, 14] *of order k*, each node is labeled with a distinct sequence of length *k* and there is an arc from one node to another if and only if there is an overlap of length *k -* 1 from the suffix of the first to the prefix of second.

Informally, a walk *p* in a graph *G* is a sequence of connected nodes by arcs. Given a sequence graph *G*, a walk *p* in *G* can spell a sequence *s*^*′*^ by concatenating the characters associated with each node of *p*. The *Graph Sequence Mapping Problem* – `GSMP` consists of finding a walk *p* in a sequence graph *G* that spells a sequence as similar as possible to a given sequence *s*.

The `GSMP`, restricted to simple sequence graph, was first addressed by Manber and Wu that propose an algorithm to find an approximated pattern in a hyper-text [16]. Akutsu proposes a polynomial time algorithm for exact mappings when hypertext is represented by a tree [15]. Park and Kim propose a polynomial time algorithm when the hypertext is represented by a directed acyclic graph [11].

One of the first articles that addresses `GSMP` in more details was written by Amir *et. al*. in the article entitled *Pattern Matching in Hypertext* [1]. Navarro improved the results of this article and detailed these improvements in the article entitled *Improved Approximate Pattern Matching on Hypertext* [8]. For the approximate mapping, Amir *et. al*. were the first authors in the literature who identified an asymmetry in the location of the changes, showing the importance of understanding whether changes happen only in the pattern, only in the hypertext or in both. Considering the asymmetry identified by Amir *et. al*., the `GSMP` allows three variants:

1. allows changes only in the pattern when mapping the pattern in hypertext;
2. allows changes only in the hypertext when mapping the pattern in hypertext;
3. allows changes in the pattern and hypertext when mapping the pattern in hypertext.

For variant 1, Amir *et. al* proposed an algorithm that runs in *O*(|*V* | +*m*·|*A*|) time which was improved by Navarro to run in *O*(*m*(|*V*| + |*A*|)) time. Here, |*V*| is the number of nodes in the graph, *A* is the number of arcs, and *m* is the length of the mapped pattern. For variants 2 and 3, Amir *et. al*. proved that the respective problems are NP-complete considering the Hamming and edit distance when the alphabet *Σ* has |*Σ*| ≥ |*V* |.

More recently, the `GSMP` restricted to simple sequence graph was addressed in the article entitled *Aligning Sequences to General Graphs in O*(|*V*| + *m* · |*A*|) *time* [12]. In this work, Rautiainen and Marschall propose an algorithm for the variant 1 that runs in *O*(|*A*|·*m*+ |*V*|·*m*.log(|*A*|·*m*)) time, which we refer to as the `GSMP` algorithm. Additionally, they also propose a more efficient version of the algorithm that returns only the distance in *O*(|*V* |+*m*·|*A*|) time, which we refer to as the `GSMP`_*d*_ algorithm. In the work entitled *On the Complexity of Sequence-to-Graph Alignment* [3], Jain *et. al* propose a correction in the algorithm proposed by Rautiainen and Marschall. They also prove that the variants 2 and 3 are NP-complete when the alphabet *Σ* has |*Σ*| ≥ 2.

The first time the `GSMP` was addressed using a De Bruijn graph as input was in the article entitled *Read Mapping on De Bruijn Graphs* [2]. In this work, Limasset *et. al*. propose the following problem, called here *De Bruijn Graph Sequence Mapping Problem* – `BSMP`: given a De Bruijn graph *G*_*k*_ and a sequence *s*, the goal is to find a path *p* in *G*_*k*_ such that the sequence *s*^*′*^ spelled by *p* have at most *d* differences between *s* and *s*^*′*^ with *d* ∈ ℕ. The `BSMP` was proved to be NP-complete considering the Hamming distance, leading to the development of a seed-and-extended heuristic by the mentioned authors. Note that for the `BSMP` it does not make sense to talk about the three variants mentioned above since there are no node repetitions.

Recently, the `BSMP` was addressed for walks in the article entitled *On the Hardness of Sequence Alignment on De Bruijn Graphs* [4]. Aluru *et. al*. proved that the problem is NP-complete when the changes occur in the graph and they proved that there is no algorithm faster than *O*(|*A*|·*m*) for De Bruijn graphs when we have changes only occur in *s* in which |*A*| is a number of arcs and *m* is the length of *s*.

Given the practical importance of the graph sequence mapping problem and the extensive use of De Bruijn graphs in the representation of sequencing data, the main focus of this work lies on a comprehensive study of the sequence mapping problem on De Bruijn graphs restricted to changes in the sequence. The main results of this study include an exact algorithm for the task of finding a walk *p* in a De Bruijn Graph *G*_*k*_ that best spells a sequence *s* under the edit distance and the development of three heuristics for the same problem which are much more efficient both in memory and time than the proposed algorithm.

This work is organized as follows: in Section 2, we describe the basic concepts we use throughout this work. In Section 3 we present the main contributions of this paper that are approaches for the `BSMP`: one adaptation of the `GSMP` algorithm and we call it here De Bruijn Sequence Mapping Tool – `BMT`, one adaptation of the `GSMP`_*d*_ algorithm and we call it here `BMT`_*d*_, one heuristic using `BMT`, two heuristics using seed-and-extend and one heuristic using `BMT`_*d*_. In Section 4 we perform experiments. Finally, in Section 5 we discuss the results and perspectives.

## 2 Preliminaries

In this section, we describe some necessary concepts such as computational definitions and problem definition that are used in this paper.

### 2.1 Sequence, distance and graphs

Let *Σ* be an **alphabet** with a finite number of characters. We denote a sequence (or string) *s* over *Σ* by *s*[1]*s*[2] … *s*[*n*] in which each character *s*[*i*] ∈ *Σ*. We say that the **length** of *s*, denoted by *s*, is *n* and that *s* is a *n***-length** sequence. We say that the sequence *s*[*i*]*s*[*i* + 1] … *s*[*j*] is a **substring** of *s* and we denote it by *s*[*i, j*]. A substring of *s* with length *k* is a *k*-length sequence and also called *k***-mer** of *s*. For 1 ≤ *j* ≤ *n* in which *n* = |*s*|, a substring *s*[1, *j*] is called a **prefix** of *s* and a substring *s*[*j, n*] is called a **suffix** of *s*.

Given five sequences *s, t, x, w, z*, we define *st* the **concatenation** of *s* and *t* and this concatenation contains all the characters of *s* followed by the characters of *t*. If *s* and *t* are *n*- and *m*-length sequences respectively, *st* is a (*n* + *m*)-length sequence. If *s* = *xw* (*x* is a prefix and *w* is a suffix of *s*) and *t* = *wv* (*w* is a prefix and *z* is a suffix of *t*), we say the substring *w* is an **overlap** of *s* and *t*.

The **Hamming distance** *d*_*h*_ of two *n*-length sequences *s* and *t* is defined as

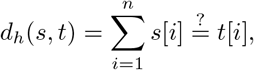

where 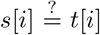 is equal to 1 if *s*[*i*] ≠ *t*[*i*], and 0 otherwise. In this context, we also say that *s* and *t* have *d*_*h*_(*s, t*) differences.

The **edit distance** is the minimum number of edit operations (insertion, deletion and substitution) required to transform one sequence onto another. Formally, the edit distance *d*_*e*_(*s, t*) between *s* and *t*, such that *n* = |*s*| and

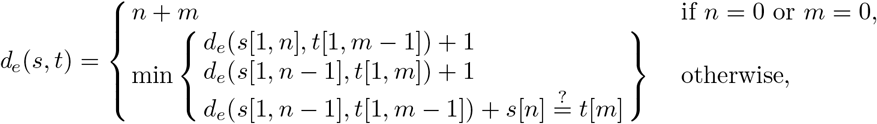

Where 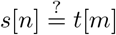 is equal to 1 if *s*[*n*] ≠ *t*[*m*], and 0 otherwise.

A **graph** is an ordered pair (*V, A*) of two sets in which *V* is a set of elements called **nodes** (of the graph) and *A* is a set of ordered pairs of nodes, called **arcs** (of the graph). Given a graph *G*, a **walk** in *G* is a sequence of nodes *p* = *v*_1_, …, *v*_*k*_, such that for each pair *v*_*i*_,*v*_*i*+1_ of nodes in *p* there is a arc (*v*_*i*_, *v*_*i*+1_) ∈ *A*. A **path** in *G* is a walk with no repeated nodes. Given a walk *p* = *v*_1_, …, *v*_*k*_, |*p*| = *k* − 1 is the **length** of *p*. For graphs with costs associated with their arcs, the **cost** of a walk *p* is the sum of the cost of all arcs of all consecutive pairs of nodes (*v*_*i*_, *v*_*i*+1_) in *p*. A **shortest path** from *v*_1_ to *v*_*k*_ is one whose cost is minimum (a path of **minimum cost**).

A **sequence graph** is a graph (*V, A*) with a sequence of characters, built on an alphabet *Σ*, associated with each of its nodes. A **simple sequence graph** is a graph in which each node is labeled with only one character. Given a set *S* = {*r*_1_, …, *r*_*m*_} of sequences and an integer *k* ≥ 2, a **De Bruijn graph** is a graph *G*_*k*_ = (*V, A*) such that:

- *V* = {*d* ∈ *Σ*^*k*^| such that *d* is a substring of length *k* (*k*-mer) of *r* ∈ *S* and *d* labels only one node};
- *A* = {(*d, d*^*′*^)| the suffix of length *k* − 1 of *d* is a prefix of *d*^*′*^}.

In this paper, informally for readability, we consider the node label as node. Given a walk *p* = *v*_1_, *v*_2_, …, *v*_*n*_ in a De Bruijn graph *G*_*k*_, the **sequence spelled** by *p* is the sequence *v*_1_*v*_2_[*k*] … *v*_*n*_[*k*], given by the concatenation of the *k*-mer *v*_1_ with the last character of each *k*-mer *v*_2_, …, *v*_*n*_. For a walk *p* = *v*_1_, *v*_2_, …, *v*_*n*_ in a simple sequence graph *G*, the sequence spelled by *p* is *v*_1_*v*_2_ … *v*_*n*_. A **mapping** of a sequence *s* onto a simple sequence graph or a De Bruijn graph *G* is a walk *p* in *G* whose editing cost between *s* and the sequence spelled by *p* is minimum.

Given the definitions above, we state the following problem for simple ssequence graphs (`GSMP`) and for De Bruijn graphs (`BSMP`) when we have changes only in the sequence, respectively:

*Problem 1 (****Graph Sequence Mapping Problem –*** `GSMP`). Given a *m*-sequence *s* and a simple sequence graph *G*, find a mapping of *s* onto *G*.

*Problem 2 (****De Bruijn Graph Sequence Mapping Problem –*** `BSMP`). Given a *m*-sequence *s* and a De Bruijn graph of order *k, G*_*k*_, find a mapping of *s* onto *G*_*k*_.

When the changes are allowed only in the sequence, we have a polynomial time algorithm proposed firstly by Amir *et. al* and Navarro [1, 8] and adapted afterward by Rautiainen and Marschall [12], that solve it. For the `BSMP`, considering paths, Limasset *et. al* [2] proved that the problem is NP-complete. More recently, for walks, Aluru *et. al* [4] proved that there is no algorithm faster than *O*(|*A*| · |*s*|) when only changes in the sequence are allowed, in which *A* is a number of arcs and |*s*| is the |*s*| -length.

Given that the results of our work are strongly based on the Rauatiainen and Marschall algorithm, we show next how it works, starting with the definition of a multilayer graph. Given a *n*-sequence *s* and a simple sequence graph *G* = (*V* = {*v*_1_, …, *v*_*m*_}, *A*), the **multilayer graph** *G*^*′*^ = (*V* ^*′*^, *A*^*′*^) is a graph obtained from *G* and *s* with weight *σ*(*e*) ∈ {0, 1} for each *e* ∈ *A*^*′*^. Informally, each layer do *G*^*′*^ is a “copy” of *G*. More precisely, the node set *V* ^*′*^ is (*V* ∪{*v*_0_}×{1, 2, …, *n*})∪{u, w} and arc set is *A*^*′*^ = *S* ∪ *T* ∪ *L* ∪ *D* ∪ *I* where

1. *S* = {(u, (*v*_*j*_, 1)) : 0 ≤ *j* ≤ *m*} with

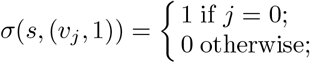
2. *T* = {((*v*_*j*_, *n*), w) : 0 ≤ *j* ≤ *m*}, with

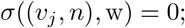
3. *L* = {((*v*_*j*_, *i*), (*v*_*h*_, *i*)) : 1 ≤ *i* ≤ *n* ∧ (*v*_*j*_, *v*_*h*_) ∈ *A*}, with

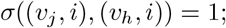
4. *D* = {((*v*_*j*_, *i*), (*v*_*h*_, *i*^*′*^)) : 1 ≤ *i < i*^*′*^ ≤ *n* ∧ *v*_*j*_ = *v*_*h*_ ∧ *i* + 1 = *i*^*′*^}, with

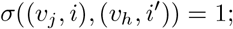
5. *I* = {((*v*_*j*_, *i* − 1), (*v*_*h*_, *i*))((*v*_0_, *i* − 1), (*v*_0_, *i*)) : *h ≠* 0, 1 *< i* ≤ *n*, (*v*_*j*_, *v*_*h*_) ∈ *A* or *j* = 0}, with

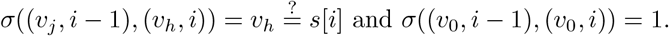

The 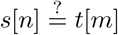 is equal to 1 if *s*[*n*] ≠ *t*[*m*], and 0 otherwise. Nodes u, w, (*v*_0_, *j*) are called *source, target* and *dummy* nodes respectively. Multilayer graph is proposed by Rautiainen and Marschall [12] to solve `GSMP` by finding a shortest path from u to w in *G*^*′*^ through a dynamic programming algorithm. Figure 1 shows an example of multilayer graph obtained from a sequence and a simple sequence graph.

**Figure 1:**
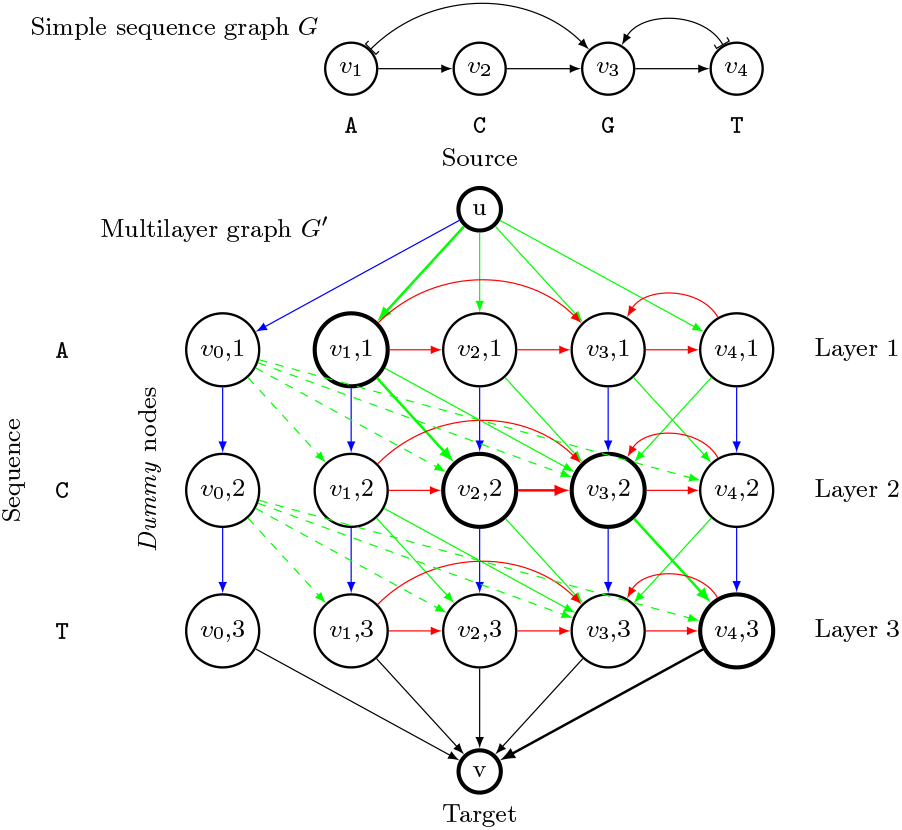
Example of mapping the sequence `s`=`ACT` in the simple sequence graph `G`. The weight of the red, blue, and green arcs are the insertion, deletion, and substitution weights, respectively. [[3, 12]].

#### Rautiainen and Marschall algorithm

Given the definition of the multilayer graph, the solution proposed by Rautiainen and Marschall to map *s* onto simple sequence graph *G* consists of building the multilayer graph *G*^*′*^ for *s* and *G* and to find a shortest path (path of minimum cost) in *G*^*′*^ from u to v. An example of a mapping based on this solution is shown in Figure 1. For this same example, a possible solution is the walk *w* = *v*_1_, *v*_2_, *v*_3_, *v*_4_ in the simple sequence graph *G*, spelled by the path *p* = *u*, (*v*_1_, 1), (*v*_2_, 2), (*v*_3_, 2), (*v*_4_, 3), *v* in *G*^*′*^. The sequence spelled by *w* corresponds to `ACGT`, whose distance to the example sequence *s* =`ACT` is

In the paper entitled by *Aligning Sequences to General Graphs in O*(|*V*| + *m*·|*A*|) *time* [12], the authors propose the algorithm detailed above that return both the mapping and the distance between the two sequences involved in *O*(|*A*|·*m* + |*V*|·*m*.log(|*A*|· *m*)) time, and we call it here the `GSMP` algorithm. In the same paper, the authors also propose a more efficient version of the algorithm that returns only the distance in *O*(|*V* | +·*m*|*A*|) time and we call it here the `GSMP`_*d*_ algorithm, in which *A* and *V* is a set of arcs and nodes, respectively and *m* is a length of mapped sequence *s*. The details of this algorithm is omitted, but its main idea is to compute the distance min *d*_*e*_(*s, s*^*′*^) over all spelled sequence *s*^*′*^ of all walks *p* in simple sequence graph *G* and for this the authors propose an algorithm that considers only two rows of the multilayer graph in each processing step, that is, for *i* = 1, …, *m* − 1, the algorithm builds rows *i* and *i* + 1, processes the two rows, deletes row *i*, advances with *i* and repeats the process until finished.

## 3 Approaches for the De Bruijn Graph Sequence Mapping Problem

In this section, we provide a detailed overview of different approaches to solve the De Bruijn Graph Sequence Mapping Problem – `BSMP`. First, we introduce an adaptation of the `GSMP` algorithm (applicable to `GSMP`_*d*_) that allow it to run with De Bruijn graphs. Then, we present three heuristics for the `BSMP`, all of which aim to find the mapping between the sequence and the graph. Two of these heuristics use the seed-and-extend strategy and one uses `GSMP` algorithm. Finally, we introduce a heuristic for the `BSMP` that only finds the distance using the `GSMP`_*d*_ algorithm.

### 3.1 An algorithm for the De Bruijn Graph Sequence Mapping Problem

The algorithm proposed here for the `BSMP` follows the same idea as the `GSMP` algorithm, but with an additional preprocessing step that converts the input De Bruijn graph into a simple sequence graph. Given a De Bruijn graph *G*_*k*_, it can be converted to a simple sequence graph following two steps: first, each node *v* of the *G*_*k*_ is splitted into *k* nodes *v*_1_ … *v*_*k*_ labeled with *v*_1_ = *v*[1], …, *v*_*k*_ = *v*[*k*]; in a second step, an arc is inserted for each pair of consecutive nodes resulting from the split of a node of *G*_*k*_, and for each pair of adjacent nodes of this graph, an arc is inserted connecting the *k*-th character of the first node with the *k*-th character of the second node. Figure 2 shows an example of this conversion and the corresponding conversion code is shown in Pseudocode 1.

**Figure 2:**
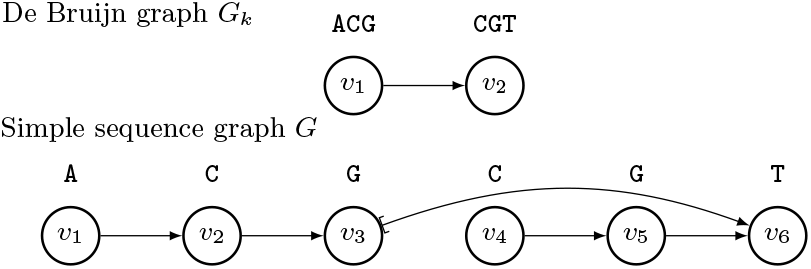
Example of converting a De bruijn graph `G`_*k*_ to a simple sequence graph `G`.

#### Converting a De Bruijn graph to a simple sequence graph works

It is easy to see that the conversion is equivalent. Given a De Bruijn graph *G*_*k*_ and the simple sequence graph *G* resulting from the conversion, note that *G* contains all *k*-mers of *G*_*k*_ splitted and correctly connected with arcs. Given *p* = *v*_1_ … *v*_*n*_ a walk in *G*_*k*_, there is a walk *qp*^*′*^ in which *p*^*′*^ = *v*1*′*, …, *vn′* in *G* and, as a result of splitted, before *v*1*′* a path *q* = *u*_1_, …, *u*_*k−*1_ with *u*_*k−*1_ = *v*1*′*, so

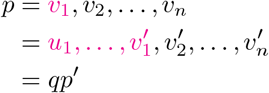

##### Pseudocode 1 convertGraph

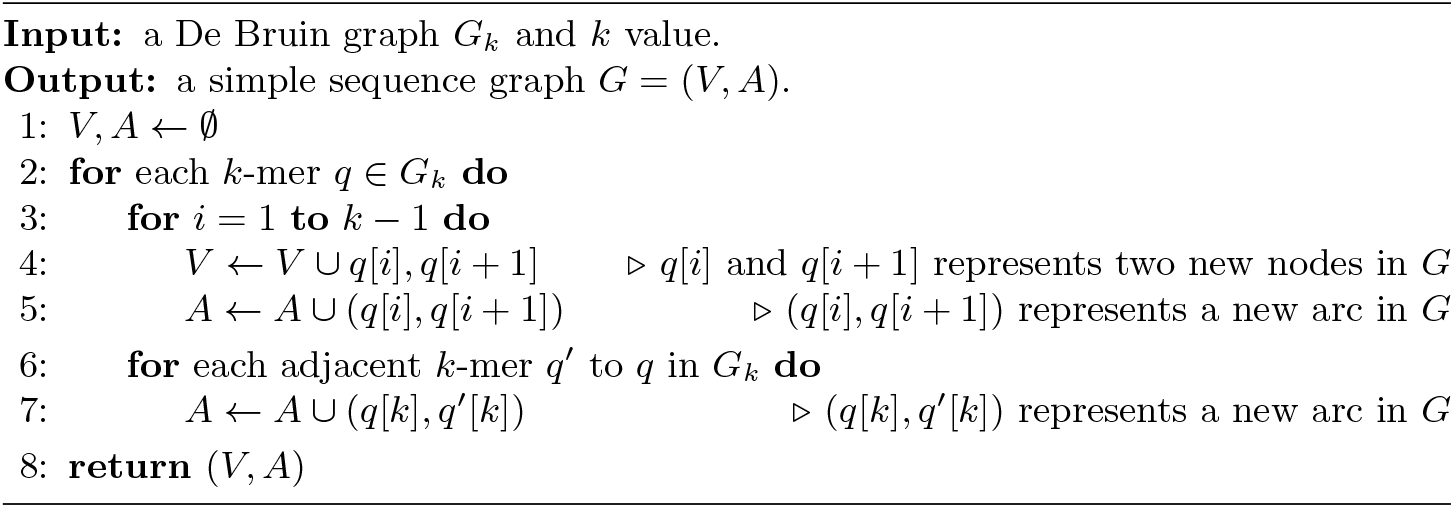

it is easy to see that the path *q* = *u*_1_, …, *v*_1_^*′*^ spells the same sequence as the node *v*_1_, and each character is spelled equivalently in *v*_2_[*k*] = *v*2*′*, …, *v*_*n*_[*k*] = *vn′*. Therefore *p* and *qp*^*′*^ spell the same sequences.

Given the simple sequence graph *G* resulting from the conversion detailed above, a solution to the `BSMP` can be achieved running the `GSMP` (or the `GSMP`_*d*_) algorithm taking *G* as input. We call this algorithm, made up of the conversion step and running of the `GSMP` (or the `GSMP`_*d*_) algorithm, the De Bruijn sequence Mapping Tool – `BMT` (or `BMT`_*d*_).

Given a sequence *s*, a De Bruijn graph *G*_*k*_ = (*V, A*), the simple sequence graph *G* = (*V* ^*′*^, *A*^*′*^) obtained from *G*_*k*_ and the multilayer graph *M* obtained from *s* and *G*, the time complexity of our approach `BMT` is

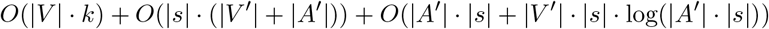

in which *O*(|*V* |·*k*) is to convert a De Bruijn *G*_*k*_ into to a simple sequence graph *G, O*(|*s*|.(|*V* ^*′*^| + |*A*^*′*^|)) is to build *M*, and *O*(|*A*^*′*^|·|*s*| + |*V* ^*′*^| |*s*·|log(|*A*^*′*^|. |*s*|)) to run `GSMP` algorithm.

The main problem of the `BMT` is in the construction of the multilayer graph. In real cases a De Bruijn graph can have thousands of *k*-mers, for example, for *k*=32 we have |*Σ*|^32^ possible *k*-mers in which *Σ* is the number of characters in the alphabet and the mapped sequences usually have lengths greater than 10000. When we build the multilayer graph, we have a copy of a De Bruijn graph converted into the simple sequence graph for each character of the sequence *s* and it is easily unfeasible to store so many graphs. For this reason, heuristics that reduce the size of the multilayer graph are welcome and may allow mapping in real cases. For our approach `BMT`_*d*_ the time complexity is

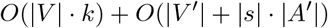

in which *O*(|*V* |·*k*) is to convert a De Bruijn *G*_*k*_ into to a simple sequence graph *G* and *O*(|*V* ^*′*^| + |*s*|·|*A*^*′*^|) to run `GSMP`_*d*_ algorithm.

`BMT`_*d*_, unlike `BMT`, is able to run real cases because the tool does not need to store the entire multilayer graph in memory, but the main problem is that we must update all the arcs when we are building a new row of the multilayer graph and that is its bottleneck in time. To resolve this question we propose a heuristic that improves your time considerably.

### 3.2 Heuristics

**Heuristic 1** Given a sequence *s* and a De Bruijn graph *G*_*k*_, the idea of our first heuristic for the `BSMP`, called here `BMT`_*h*1_, is to anchor some *k*-mers of the sequence *s* into the graph and use the `BMT` to map only the substrings of *s* that could not be anchored. In more details, the first step of the `BMT`_*h*1_ is to look for all the *k*-mers in *s* that are present in *G*_*k*_, that we call **anchors**. Note that, after this step, we can have several substrings of *s* built by *k*-mers of the sequence that are not in *G*_*k*_, that we call **gaps**. For each one of these gaps bordered by two anchors, the heuristic use the `BMT` to find a walk that, starting at the anchor immediately to the left of the gap and ending at anchor immediately to the right of the gap, best fill it. Figure 3 and 4 shows an example of how `BMT`_*h*1_ proceeds, in the first step (Figure 3) we determine all anchors and in the second step (Figure 4) we determine, with `BMT`, the best walks between two anchors The wavy lines are the walks (found with the `BMT` algorithm) that best fill each gap (starting at the first anchor and ending at the second anchor).

**Figure 3:**
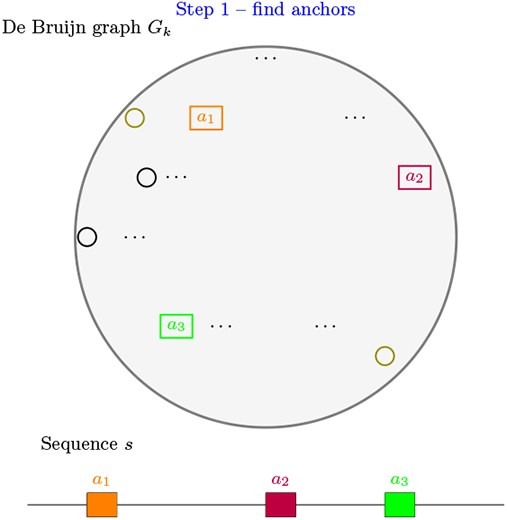
First step – each square is an anchor found in `G`_k_.

**Figure 4:**
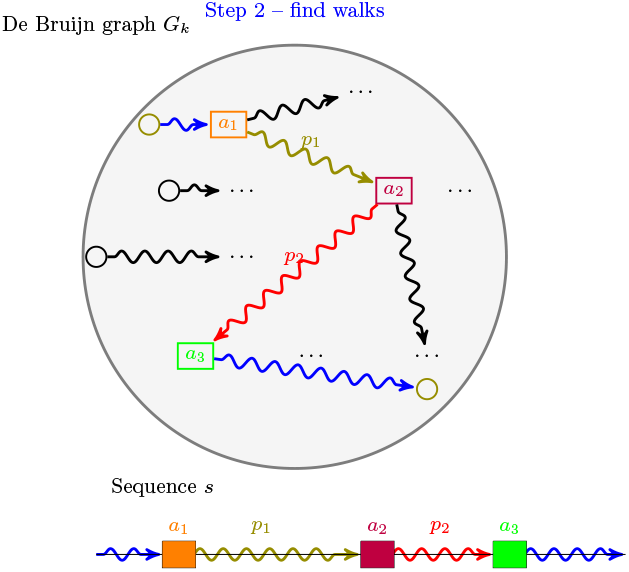
Second step – The wavy lines between the anchors are the best walks found between them with `BMT` algorithm.

We show in Pseudocode 2 the steps that `BMT`_*h*1_ follows and we show in Pseudocode 3 the steps that **findAnchors** follows. **FindAnchors** returns a sequence of anchors *H*, and for each consecutive anchors *a, a*^*′*^ ∈ *H* the `BMT`_*h*1_ uses `BMT` to find the best walk from *a* to *a*^*′*^.

Experimental results show that it is costly to consider the entire De Bruijn graph in each call of `BMT` in Heuristic 2. For this reason, `BMT`_*h*1_ considers only a subset of *V* (*G*_*k*_) in the search for a best walk between two consecutive anchors: considering that the gap has size *N*, only nodes that are 0.5 *N* distant, in terms of their arcs, from the source and target nodes are considered. Considering this constraint, the heuristic is able to process larger instances, but the resulting walk may not be able to cover the entire sequence since `BMT` may not find a walk from the source and the target nodes. In this case, we consider as the final answer the longest mapping.

#### Pseudocode 2 BMT_*h*1_

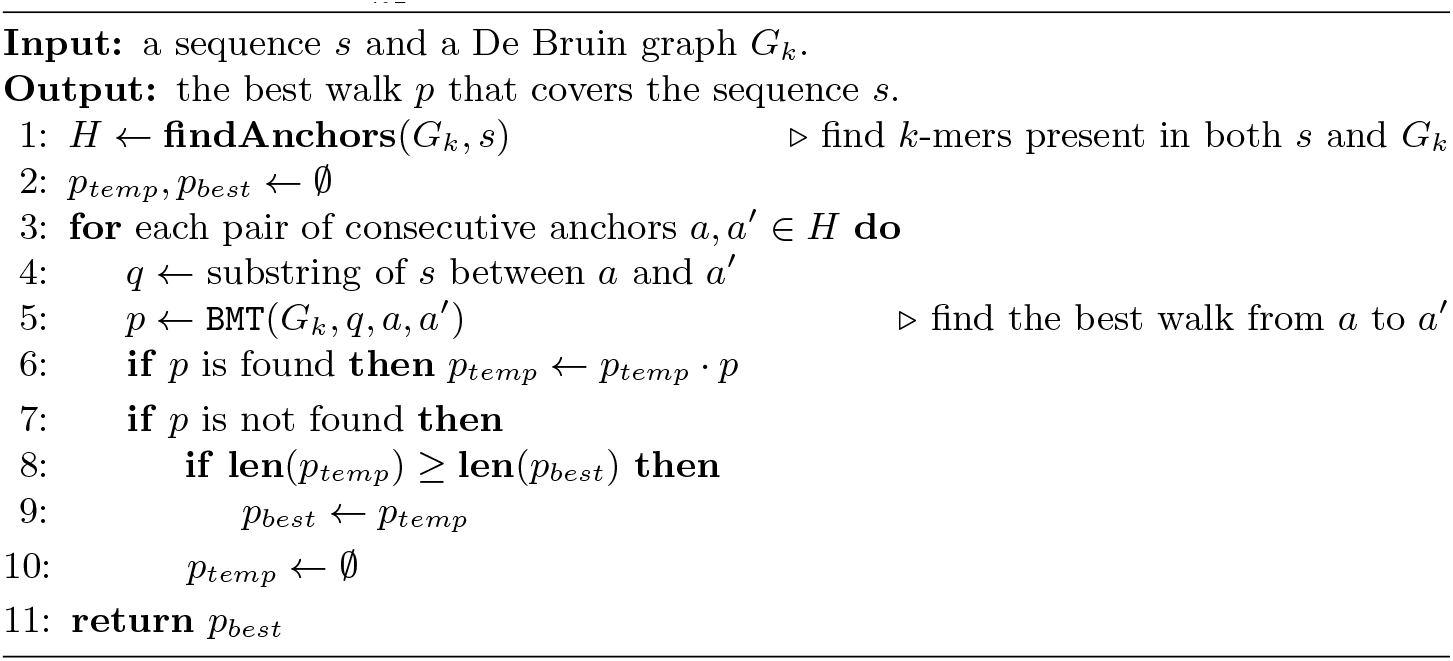

#### Pseudocode 3 findAnchors

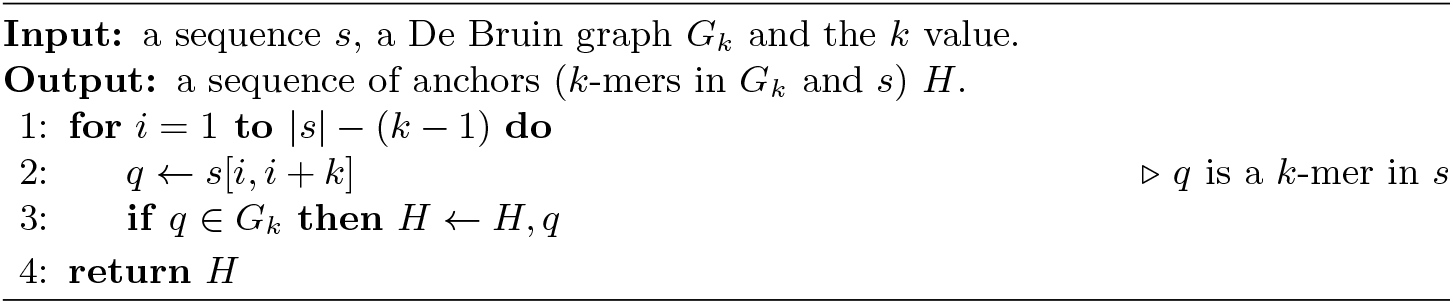

The main advantage of `BMT`_*h*1_ over `BMT` is the amount of memory used. For the `BMT` we have to build a multilayer graph considering the whole sequence and the whole De Bruijn graph, whereas for the `BMT`_*h*1_ we build the multilayer graph for the substring of the gaps and consider only a subgraph of *G*_*k*_. For the `BMT`_*h*1_ we can find the anchors in constant time and it takes *O*(|*s*| − *k*) to find all the anchors of *s*. Given *N* anchors found, the time complexity is *O*(*N* (|*A*^*′*^| |*s*| + |*V* ^*′*^| |*s*|.log(|*A*^*′*^|·|*s*|))).

We also developed the 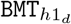 using the same idea as `BMT`_*h*1_. The 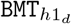 anchor some *k*-mers of the sequence *s* into the graph and use the `BMT`_*d*_ to find the distance of the best walk. In more details, the first step of the 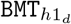 is to look for all the anchors (*k*-mers) in *s* that are present in *G*_*k*_. Note that, after this step, we can also have several gaps. For each one of these gaps bordered by two anchors, the heuristic use the `BMT`_*d*_ to find the distance of the best walk, starting at the anchor immediately to the left of the gap and ending at anchor immediately to the right of the gap. Here 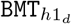 also consider only a subset of *V* (*G*_*k*_) in the search for distance of the best walk between two consecutive anchors and the 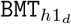 consider as the final answer the approximated distance of the longest mapping. The `BMT`_*d*_ is capable of processing large inputs, but it takes a long time to run and the main advantage of 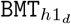 over `BMT`_*d*_ is in runtime because we run `BMT`_*d*_ only for gaps and it is advantageous. For the 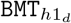we can find the anchors in constant time and it takes *O*(|*s*|− *k*) to find all the anchors of *s*. Given *N* anchors found, the time complexity is *O*(*N* ·((|*V* |·*k*)+(|*V*^*′*.^|+|*s*||*A*^*′*^|)).

In order to improve the performance of `BMT`_*h*1_ and 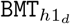, we develop two versions of it called 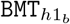 and 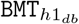 for `BMT`_*h*1_ and 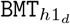 respectively. In this new versions, we build De Bruijn graphs with the Bifrost – a tool that allows to build De Bruijn graphs efficiently and run algorithms like Breadth-First Search and dijkstra on these graphs [9]. A relevant point of using Bifrost in 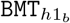 and 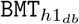 is that Bifrost has its own way of efficiently creating the De Bruijn graph and its own interpretation of *k*-mers, in general, the amount of anchors between our tools with and without Bifrost can vary and this has an impact on the final answer. We can observe this impact in tests when looking at the difference between the responses of 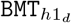 and 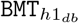, and between `BMT`_*h*1_ and 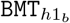.

**Heuristic 2** Another way to perform the mapping of a sequence in a De Bruijn graph is using the seed-and-extend approach, which consists of, given a set of seeds (substrings of) in the query sequence, to extend each of them in both directions (left and right) while comparing their subsequent characters with the characters of a subject sequence. These extensions proceed until a number of differences be found in this comparison process. This idea is used by us in the development of a new heuristic to the `BSMP`, called here `BMT`_*h*2_, that consists of the following steps: (1) find all anchors (seeds) in the sequence *s* (subject sequence); (2) try to extend all the anchors to the right comparing the subsequent characters with the characters (query sequence) spelled by a walk in the De Bruijn graph starting in the anchor node until one difference be found; (3) return the walk that induces the largest extension of *s*. In order to allow substitutions in the sequence *s*, for each anchor to be extended, we evaluate the possibility of changing its last character and, with this, to find a better walk in the graph starting on a new modified anchor node. If during the extension we reach another anchor, we continue the extension and this new anchor is part of the solution. Considering that we can have several extensions, the resulting walk may not be able to cover the entire sequence since `BMT`_*h*2_ may not find a walk during the extension. In this case, we consider as the final answer the longest mapping.

Figure 5 and 6 shows an example of how `BMT`_*h*2_ proceeds, in the first step (Figure 5) we determine all anchors and in the second and third step (Figure 6) we determine, with extend, the best walk between two anchors and return it. The wavy lines are the walks, and the green walk is the best walk (starting at the new modified anchor node and ending at the second anchor). For the `BMT`_*h*2_ we can find the anchors in constant time and it takes *O*(|*s*| − *k*) to find all the anchors of *s*. Given *N* anchors found and an alphabet |*Σ*| = *M*, the time complexity is *O*(|*N*| · (*M*·|*A*|)) to extend all anchors in which |*A*| is the number of arcs in the De Bruijn graph.

**Figure 5:**
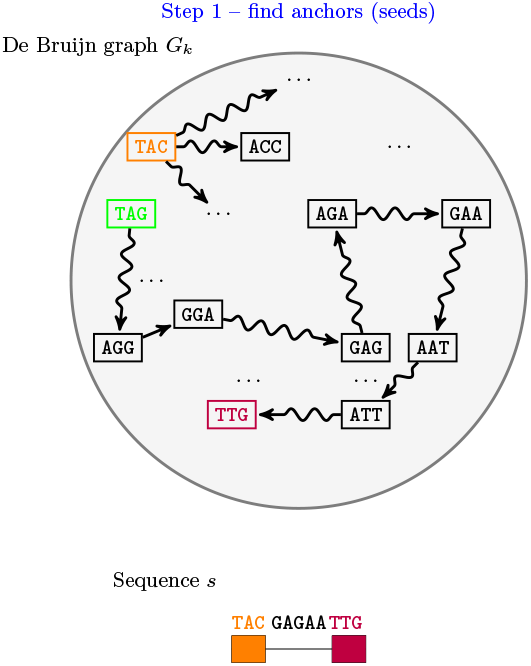
Step 1 – square orange and purple are anchors found in `G`_k_.

**Figure 6:**
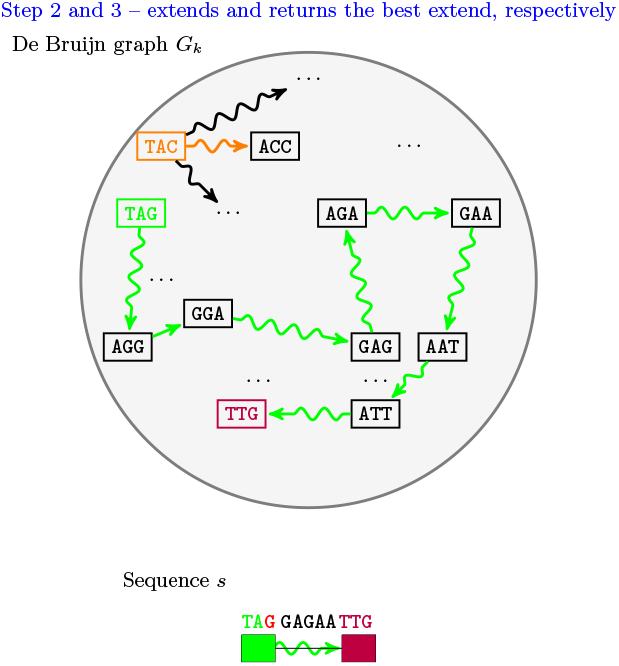
Step 2 – extends by changing the last character, for this example, it is best to change (in `s`) `TAC` to `TAG` and extend from the new anchor. Step 3 – return the best extend.

#### Heuristic 3

In order to improve the `BMT`_*h*2_ coverage, we develop a new heuristic that allows substitution and deletion in the mapped sequence, called here `BMT`_*h*3_, that consists the same steps of the `BMT`_*h*2_, but if during the extension step we fail to extend for any walks, we remove *k* − 1 characters from the anchor and repeat the extension process. For the `BMT`_*h*3_ we can find the anchors in constant time and it takes *O*(|*s*| − *k*) to find all the anchors of *s*. Given *N* anchors found and an alphabet |*Σ*| = *M*, the time complexity is *O*(|*N* |.(*M*.(|*s*| − (*k* − 1)). |*A*|)) to extend all anchors in which |*A*| is the number of arcs in the De Bruijn graph and |*s*| − (*k* − 1) is a number of deletions.

Note that in heuristic 2 we only perform substitution operations and in heuristic 3 we perform substitution and deletion operations in the sequence. All implementations are available on github: github.com/LucasBarbosaRocha.

## 4 Experiments and Results

In order to evaluate our approaches, the accuracy and performance of the proposed heuristics were assessed by a number of experiments on random(ou quasirandom) data. All the experiments were performed on a computer with processor Intel(R) Xeon(R) CPU E5-2620 2.00GHz with 24 CPUs and 62 GB of RAM. Our dataset includes more than 20 De Bruijn Graphs, build on short reads (100bp on average) of DNA sequences of Escherichia coli (set *A*) and Escherichia fergusonii (set *B*) organisms and more than 100 long reads (5000 bp on average) of Escherichia coli (set *C*) organisms to be mapped in the graph. All of these sequences were obtained from the GenBank^1^ database of the NCBI.

In Section 4.1, we compare the accuracy and runtime of the heuristics `BMT`_*h*1_ and 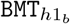 and we used the `BMT`_*d*_ to get the exact distance since `BMT` is not able of processing large numbers of *k*-mers. In Section 4.2, we compare all the four in terms of accuracy and runtime using again the `BMT`_*d*_ to get the exact distance. Finally, In Section 4.3, we compare the accuracy and runtime of the exact solution `BMT`_*d*_ and the heuristics 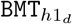 and 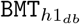.

To calculate the average of distances of the implemented approaches, we add up all the distances obtained by the heuristics `BMT`_*d*_, 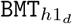 and 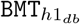 and divide this summation by the number of tests performed. For the `BMT`_*h*1_, 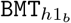, `BMT`_*h*2_ and `BMT`_*h*3_ heuristics, we take the sequences returned by the heuristics, calculate the edit distance between each of them and the original mapped sequence, add uo these distances and divide by the number of tests. To verify the accuracy of the results, we checked in the next sections for each row on average distance tables the difference between the exact distance of the algorithm `BMT`_*d*_ and that one of each heuristic.

### 4.1 Tests to obtain mapping -heuristic 1

To run the tests of this section, we use the De Bruijn graph with varying number of *k*-mers built by Set *A* and we take sequences of length between 2381 and 6225 from Set *C*. The heuristics return the string spelled by the best walk found and we calculate the edit distance between the original string and the spelled string and compare it to the exact distance returned by the `BMT`_*d*_. In these tests, the number of anchors lies between 0 and 2015 and we can see how using the Bifrost tool on 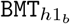 brought the distance closer to `BMT`_*d*_ than `BMT`_*h*1_.

The average distances and running runtime obtained in this round of tests are shown on Table 1 and Table 2, respectively. The charts for the calculate average distances and running time are shown on Figure 7 and Figure 8, respectively. Different from our heuristics, the `BMT` was not able to manage the vast majority of test cases. In our analyses, `BMT` managed to process small test cases for long reads with a length close to 1000 and a De Bruijn graph with approximately 2000 *k*-mers. We were able to use `BMT` and 3 out of 100 tests, and this fact attests the applicability of our approaches. We can see in Table 1 that the heuristics are able to perform the mapping in relation to `BMT` and we can see in Table 2 that the distance obtained by the heuristics are not so far away from the exact cost obtained by `BMT`_*d*_.

**Table 1:** Average distances and accuracy: tests with length of long reads between 2381 and 6225.

| k | Qty. $k$-mers in $G_k$ | Average distances | | | Accuracy | |
| --- | --- | --- | --- | --- | --- | --- |
|  |  | BMT<sub>d</sub> | BMT<sub>h1b</sub> | BMT<sub>h1</sub> | BMT<sub>h1b</sub> | BMT<sub>h1</sub> |
| 10 | 78661 | 1746,4 | 3046,8 | 5623,0 | 1,7 | 3,2 |
| 15 | 86214 | 2416,3 | 5197,5 | 6255,8 | 2,2 | 2,6 |

**Table 2:** Average time (seconds): tests with length of long reads between 2381 and 6225.

| k | Qty. $k$-mers in $G_k$ | BMT<sub>h1</sub> | BMT<sub>h1b</sub> |
| --- | --- | --- | --- |
| 10 | 78661 | 2250,0 | 1125,0 |
| 15 | 86214 | 3160,3 | 1365,0 |

**Figure 7:**
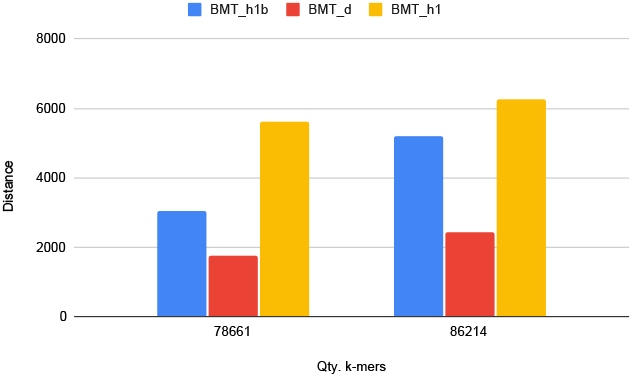
Average distances: chart for Table 1.

**Figure 8:**
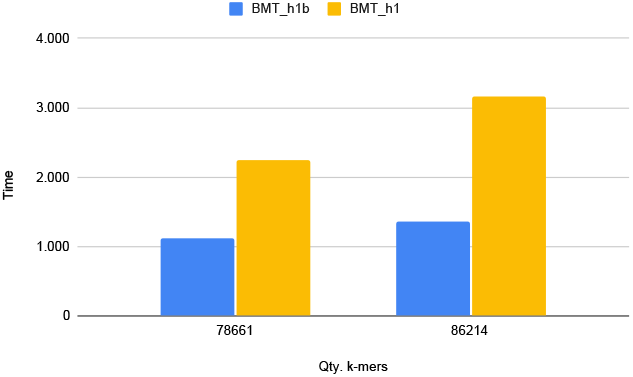
Average time (seconds): chart for Table 2.

### 4.2 Tests to obtain mapping -all heuristics

To run the tests of this section, we use De Bruijn graphs with a varying amount of *k*-mers built by Set *B* and we take sequences of length between 2381 and 6225 from Set *C*. The calculated average distances are shown on Table 3 and 4 and the respective chars for each table shown on Figure 9 and Figure 10. We can see that 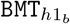 is the best heuristic in time and quality and is our most powerful version because its graph construction tool (Bifrost) is capable of creating much larger graphs. `BMT`_*h*3_ is a heuristic that, despite being simple, manages to be the second best and deliver results in an accessible time. Another important detail is that the number of anchors is relevant for the good execution of the heuristics `BMT`_*h*1_ and 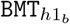 and the greater the number of anchors, the better the answer will be and therefore we will process less of the sequence. For heuristics `BMT`_*h*2_ and `BMT`_*h*3_, anchors are not used, but it is important to note that a sequence with many anchors is a sequence very similar to the graph and this indicates that the `BMT`_*h*2_ and `BMT`_*h*3_ heuristics will obtain better answers.

**Table 3:**
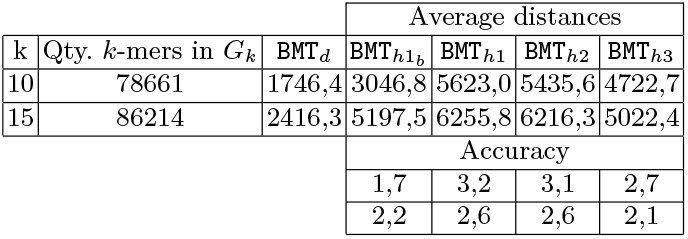
Average distances and accuracy: tests with length of long reads between 2381 and 6225.

**Table 4:** Average time (seconds): tests with length of long reads between 2381 and 6225.

| k | Qty. $k$-mers in $G_k$ | $BMT_{h1b}$ | $BMT_{h1}$ | $BMT_{h2}$ | $BMT_{h3}$ |
| --- | --- | --- | --- | --- | --- |
| 10 | 78661 | 1125,0 | 2250,0 | 1508,4 | 2160,6 |
| 15 | 86214 | 1365,0 | 3160,3 | 1882,5 | 4560,0 |

**Figure 9:**
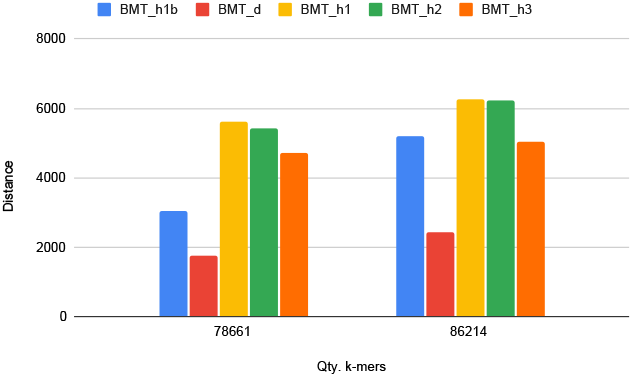
Average distances: chart for Table 3.

**Figure 10:**
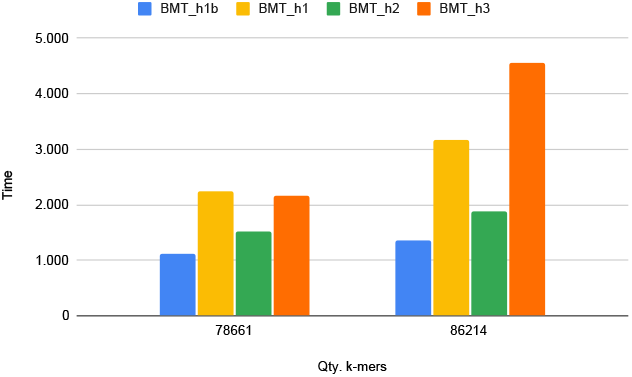
Average time (seconds): chart for Table 4.

We perform another tests with sequences of length between 1000 and 10000 from Set *C* and we use De Bruijn graphs built by Set *A*. The calculated average distances are shown on Table 5 and the average running times of each heuristic shown on Table 6, with the respective charts for each table shown of Figure 11 and 12. In these new tests we get several sequences with length between 1000 and 10000; and the value of *k* varying between 5, 10 and 20. In these new tests we can verify a more interesting competitiveness in terms of time and quality. The heuristics were very close in values and heuristics `BMT`_*h*2_ and `BMT`_*h*3_ showed their efficiency in achieving results close to the other heuristics.

**Table 5:**
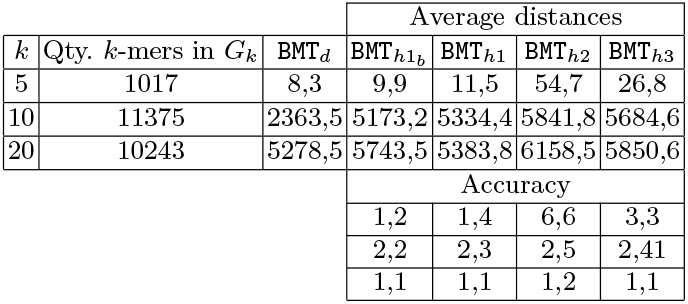
Average distances and accuracy: tests with length of long reads between 1000 and 10000.

**Table 6:**
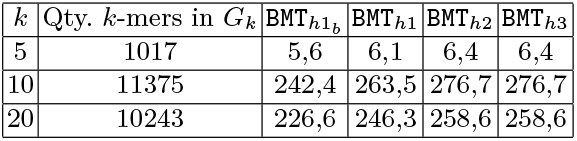
Average time (seconds): tests with length of long reads between 1000 and 10000.

**Figure 11:**
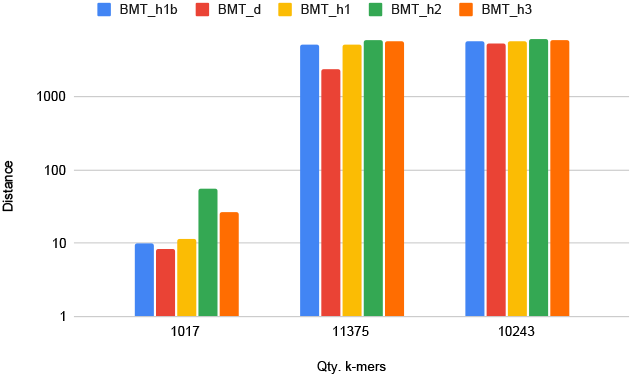
Average distances: chart for Table 5.

**Figure 12:**
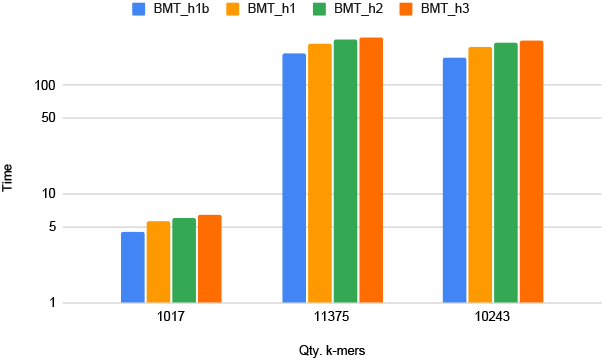
Average time (seconds): chart for Table 6.

Here the number of anchors is between 0 and 9234. For the small k 5 we have the largest number of anchors and this gives us a very mapped sequence. For k 10 we have the amount of anchors being around 50% of the sequence length and this gives us a partially mapped sequence. For k 20 we have a special case that consists of having a few anchors and the result is very close to the `BMT`_*d*_ and this happens because our heuristics return a walk *p* that spells a small sequence and when calculating the accuracy with the original sequence we have a high edit distance and this is close to the distance returned by `BMT`_*d*_.

### 4.3 Tests to obtain only distances -heuristic 1

A relevant test is to want to find only the mapping distance (and not the mapping) and in this section we perform this test. To run the tests of this section, we use De Bruijn graphs with a varying amount of *k*-mers built by Set *A* and we take sequences of length between 2381 and 6225 from Set *C*. The calculated average distances are shown on Table 7 and the average running times of each heuristic are shown on Table 8, with the respective charts for each table shown on Figure 13 and 14. We can see that for the version that only looks for the distance, it is possible to have a great time gain with heuristics compared to `BMT`_*d*_ because we take advantage of anchors to not process the entire sequence, on the other hand, we have a loss of accuracy with the heuristics. The anchors in these tests increase as the number of *k*-mers in the graph increases, with their number varying between 23 and 6000 anchors.

**Table 7:** Average distances and accuracy: tests with length of long reads between 2381 and 6225 and the `k` = 10.

| Qty. $k$-mers in $G_k$ | Average distances | | | Accuracy | |
| --- | --- | --- | --- | --- | --- |
|  | BMT<sub>d</sub> | BMT<sub>h1d</sub> | BMT<sub>h1db</sub> | BMT<sub>h1d</sub> | BMT<sub>h1db</sub> |
| 4577 | 1537,5 | 3147,0 | 3181,0 | 2,1 | 2,1 |
| 9119 | 1453,3 | 3184,3 | 3098,3 | 2,2 | 2,2 |
| 11375 | 1427,0 | 3142,3 | 3028,0 | 2,2 | 2,2 |
| 22485 | 1201,7 | 3111,3 | 2835,0 | 2,6 | 2,4 |
| 43842 | 995,3 | 2850,5 | 2558,0 | 2,8 | 2,6 |
| 84683 | 662,7 | 1663,3 | 1550,0 | 2,5 | 2,4 |
| 557522 | 314,5 | 552,3 | 477,3 | 1,7 | 1,5 |

**Table 8:** Average time (seconds): tests with length of long reads between 2381 and 6225 and the `k` = 10.

| Qty. $k$-mers in $G_k$ | BMT<sub>d</sub> | BMT<sub>h1d</sub> | BMT<sub>h1db</sub> |
| --- | --- | --- | --- |
| 4577 | 73,7 | 5,0 | 3,5 |
| 9119 | 222,5 | 7,7 | 5,7 |
| 11375 | 370,0 | 32,5 | 24,7 |
| 22485 | 1632,3 | 24,0 | 13,3 |
| 43842 | 13324,0 | 261,7 | 177,7 |
| 84683 | 89152,0 | 66,0 | 41,7 |
| 557522 | 151787,3 | 45,0 | 29,7 |

**Figure 13:**
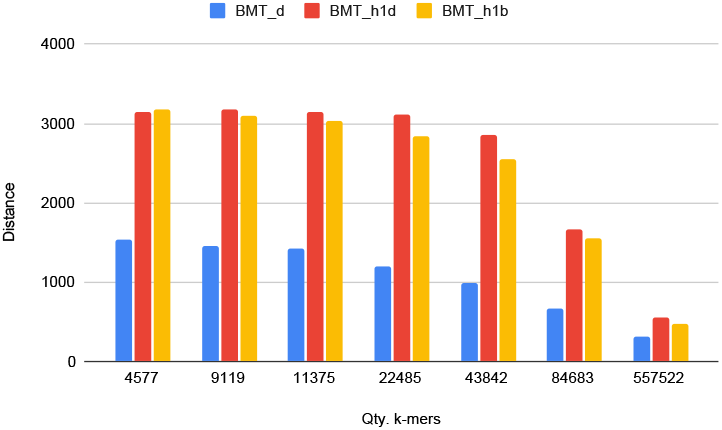
Average distances: chart for Table 7.

**Figure 14:**
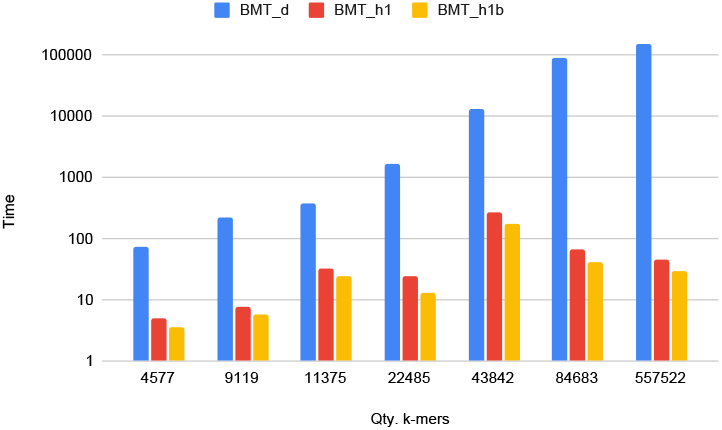
Average time (seconds): chart for Table 8.

## 5 Discussion, perspectives and conclusions

This work deals with two versions of De Bruijn Graph Sequence Problem – `BSMP`: one aims to find a minimum cost mapping between a sequence and a Bruijn graph, and the other aims to find the distance between them. For each version, we develop exact algorithms that is a simple adaptations of Rautiainen and Marschall [12] algorithm for simple sequence graphs. The developed algorithms are De Bruijn Mapping Tool – `BMT` that returns the mapping and `BMT`_*d*_ that returns only the distance between them. However, `BMT` is not capable of working with large graphs because it requires a lot of memory and `BMT`_*d*_ takes a lot of running time. Therefore, we also design heuristics that overcome these difficulties and find a low-cost mapping and an approximate distance.

`BMT` can not handle sequences longer than 1000 elements or graphs with more than 2000 10-mers. However, our best heuristic for finding the mapping of approximate distance, `BMT`_*h*1_*b* can handle graphs with 86214 15-mers and sequences up to 10000 elements in less than 22 minutes.

`BMT`_*d*_ can handle sequences with up to 7000 elements and graphs with with up 560,000 10-mers, but it takes more than 40 hours. On the other hand, our best heuristic for the approximated distance, 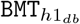, spends only 29 seconds.

From this work, several variations should be considered to improve the performance and quality of practical solutions.

Firstly, concerning the graph mapping problem, we not only propose the 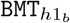 heuristic but also introduce two other heuristics, `BMT`_*h*2_ and `BMT`_*h*3_, that employ the seed-and-extend strategy. Although `BMT`_*h*2_ and `BMT`_*h*3_ exhibit lower performance than 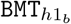, the concepts developed in these heuristics can be incorporated into 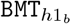, resulting in a new and more efficient heuristic version.

Hirschberg [5] reduces the quadratic space used to find an alignment for a pair of sequences using linear space by using the divide-and-conquer paradigm. Therefore, secondly, in the context of the mapping version, these ideas could be adapted to reduce the space and extend the scope of this work.

Finally, another promising approach to enhance the practical results of both versions is the parallelization of the algorithms. This approach can significantly reduce the execution time and increase the scalability of the algorithms.

## Footnotes

1 https://www.ncbi.nlm.nih.gov/genbank/

